# A two-step model of autophagy: autophagosome formation, degradation and net turnover

**DOI:** 10.1101/2020.09.29.318386

**Authors:** Ainhoa Plaza-Zabala, Virginia Sierra-Torre, Amanda Sierra

## Abstract

Autophagy is a complex process that encompasses the enclosure of cytoplasmic debris or dysfunctional organelles in membranous vesicles, the autophagosomes, for their elimination in the lysosomes. A gold-standard method to assess its induction is the analysis of the autophagic flux using as a surrogate the expression of the microtubule-associated light chain protein 3 conjugated to phosphatidylethanolamine (LC3-II) by Western blot, in the presence of lysosomal inhibitors. Therefore, the current definition of autophagy flux actually puts the focus on the degradation stage of autophagy. In contrast, the most important autophagy controlling genes that have been identified in the last few years in fact target early stages of autophagosome formation. From a biological standpoint is therefore conceivable that autophagosome formation and degradation are independently regulated and we argue that both stages need to be systematically analyzed. Here, we propose a simple two-step model to understand changes in autophagosome formation and degradation using data from conventional LC3-II Western blot, and test it using two models of autophagy modulation in cultured microglia, the brain macrophages: rapamycin and the ULK1/2 inhibitor, MRT68921. Our model provides a comprehensive understanding of the autophagy process and will help to unravel the effect of genetic, pharmacological, and environmental manipulations on both the formation and degradation of autophagosomes.

## Introduction

Autophagy is a complex phenomenon dedicated to eliminate intracellular debris, from protein aggregates to dysfunctional organelles, and is thus essential to maintain cell fitness (Plaza-Zabala et al., 2017; Levine and Kroemer, 2019). Assessing autophagy is complicated and current guidelines recommend using several complementary methods (Klionsky et al., 2016). Nonetheless, the gold standard remains the analysis of the autophagic flux using LC3 (microtubule-associated light chain protein 3). During autophagy, cytosolic LC3 (LC3-I) is conjugated to phosphatidylethanolamine and recruited to the nascent phagophore membranes (LC3-II). The phagophore then encloses cytosolic material or organelles forming a double-membrane autophagosome, which is then redirected towards the lysosome for its enzymatic degradation. The autophagic flux is calculated as the differential amount of LC3-II in the presence/absence of lysosomal inhibitors, such as bafilomycin or chloroquine, among others. Thus, the autophagic flux as is currently defined reflects the late stages of autophagy, i.e., lysosomal degradation, although as we will show here it contains information about both autophagosome formation and degradation (**Figure 1A**). Importantly, formation and degradation are regulated by concerted but independent mechanisms: most autophagic-regulatory genes are involved in the early stages of autophagy, as is the case of the ATG family, encoding proteins that are mainly involved in autophagosome formation and maturation (Plaza-Zabala et al., 2017; Mercer et al., 2018). In contrast, autophagosome degradation largely depends on lysosomal proteins and enzymes (**Figure 1B**). Therefore, both early and late stages of autophagy should be systematically analyzed to understand the autophagosome turnover in any given condition.

**Figure 1.**
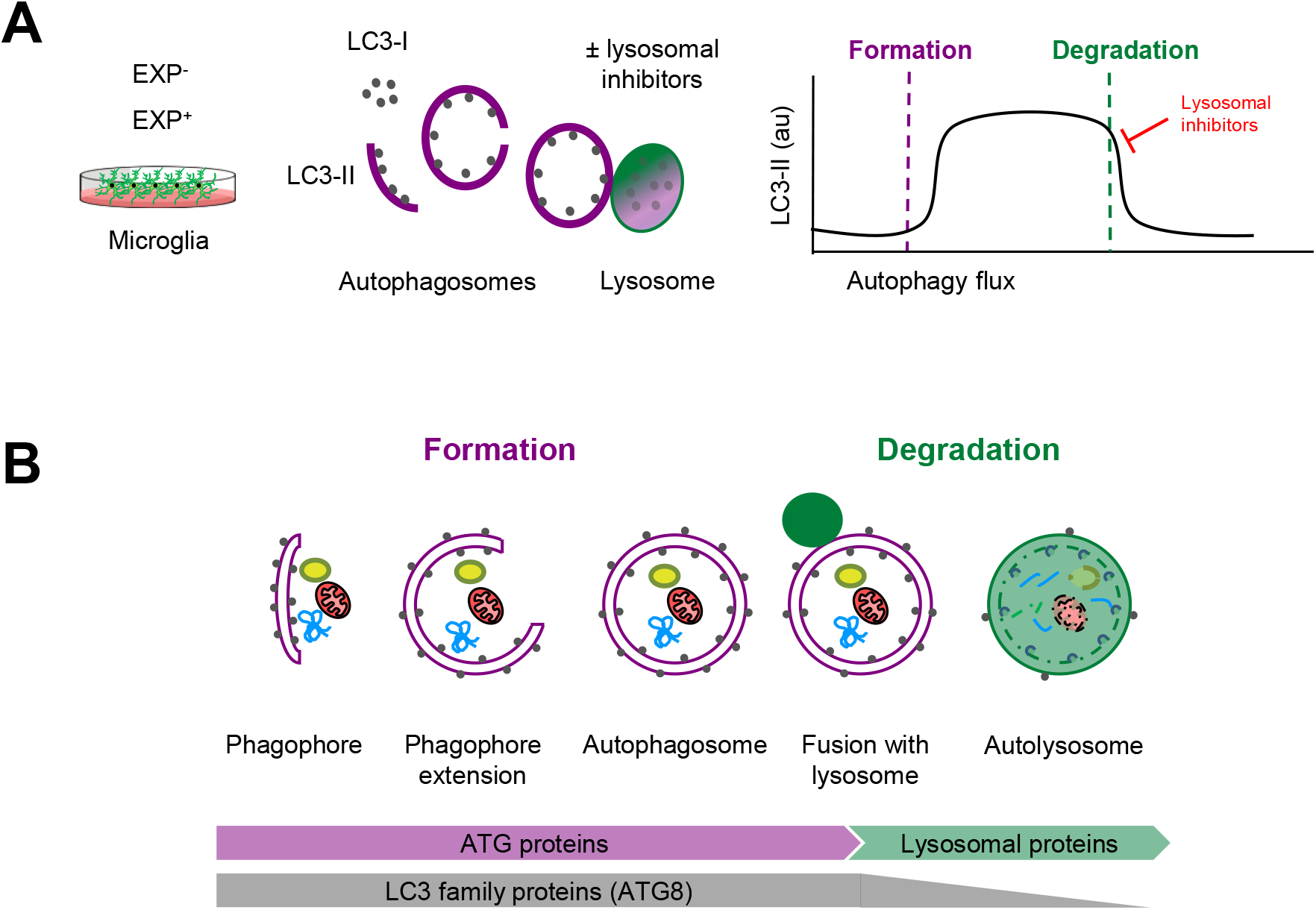
Estimation of autophagy flux variations using LC3 turnover assay. **A**, Total protein homogenates obtained from microglia under control (EXP^−^) and experimental conditions (EXP^+^) are analyzed by Western Blot to evaluate LC3 levels in the presence and absence of lysosomal inhibitors. When autophagy is activated, LC3-I (soluble form) is lipidated to the phophatidylethanolamine of the nascent phagophore forming LC3-II (membrane-bound form). LC3-II accumulates along the extension of the autophagic vacuoles as it closes and is used as an estimate of the number of autophagosomes. Upon fusion with lysosomes, LC3-II levels decrease due to the degradation of the inner autophagosomal membrane simultaneously with the luminal cargo. In the presence of lysosomal inhibitors, no degradation occurs and LC3-II levels are maintained. The subtraction of LC3-II quantities in the presence and absence of lysosomal inhibitors provides an estimate of the autophagosomes that have been degraded during the experimental period of time. **B**, Early stages of autophagy, which lead to the *de novo* formation of autophagosomes, are mainly regulated by ATG proteins. The LC3 family of proteins (ATG8) participate in the formation of autophagosomes and progressively disappear after lysosomal fusion and cargo degradation in autolysosomes. Late stages of autophagy depend on the functionality of lysosomal proteins and enzymes.

### Modeling autophagosome formation and degradation

Here we propose a simple two-step model to analyze autophagy, in which the net number of autophagosomes (i.e., the autophagosome pool) at any given time is treated as a black box to which there is an input (formation) and an output (degradation)(**Figure 2A**). The formation phase encompasses phagophore formation, cargo sequestration, and autophagosome closure, and the degradation phase summarizes the lysosomal fusion and the enzymatic degradation of the autophagosome contents. Nonetheless, the precise definition of formation/degradation in each experimental setup depends on the physiological process blocked by the particular lysosomal inhibitor used: fusion inhibitors, such as vinblastine, which blocks transport of autophagosomes by microtubules; protease inhibitors, such as E64d and leupeptin; or proton pump inhibitors, such as bafilomycin. This conceptual frame can be easily modeled by a differential equation in which the change in the size of the autophagosome (APh) pool over time (t) or after a stimulus depends on the number of autophagosomes in the steady-state (ss) plus the number of autophagosomes formed minus the autophagosomes degraded in a certain period of time:

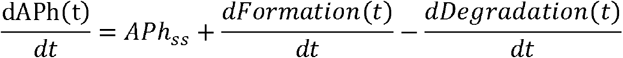

The difference between formation and degradation is the net autophagic turnover, which is a measure of the relative velocity of autophagosome formation versus degradation. A given stimulus could act both on the formation and/or the degradation, maintaining the size of the APh pool and resulting in a constant net turnover ratio (**Figure 2A1**). However, under some conditions, the regulation of the formation and degradation of autophagosomes may be dissociated: an increased degradation would decrease the size of the APh pool and increase the net turnover (**Figure 2A2**); and an increased formation would increase the size of the APh pool and decrease the net turnover (**Figure 2A3**). Thus, we propose that to understand the complexity of the biology underneath the autophagosome turnover we need to analyze separately formation, degradation, and the net autophagic turnover.

**Figure 2.**
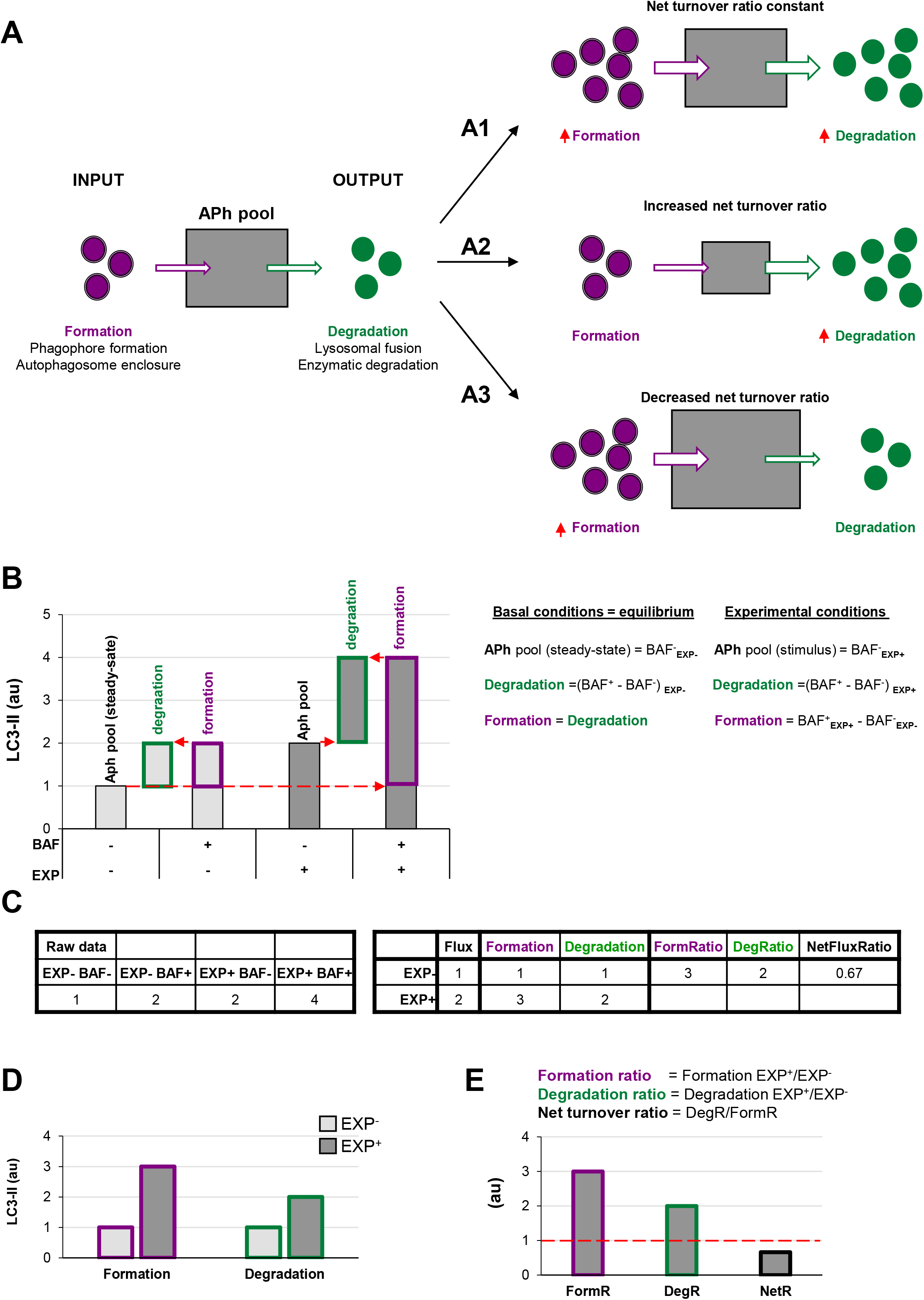
A two-step model of autophagy to analyze formation and degradation of autophagosomes. **A**, the model represents the autophagosomes as a box with an input (autophagosome formation) and an output (autophagosome degradation) that determines the autophagosome net turnover. **A1-A3** represent different possible scenarios with no changes (**A1**), an increase (**A2**) and a decrease (**A3**) in the autophagosome net turnover. **B**, graph the amount of LC3-II (au, arbitrary units) in two experimental conditions representing (EXP^−^ and EXP^+^) in the presence or absence of the lysosomal inhibitor bafilomycin (BAF^−^ and BAF^+^), and the formulas used to calculate formation, degradation and net turnover. The dotted red arrows mark the LC3-II raw data values used to calculate the formation and degradation rates and ratios. **C**, Simulated raw LC3-II data (au) (left) used to calculate the formation and degradation rates and ratios (right) used in the graphs shown in **B, D**, and **E**. **D**, **E** graphs representing the rate of change of formation, degradation and net turnover between the two experimental conditions.

This analysis can be performed using the data available in conventional LC3 assays by Western blot. In this type of analysis, the cells/tissue from two experimental conditions (EXP^−^ and EXP^+^) are incubated with lysosomal inhibitors such as bafilomycin (BAF^−^ and BAF^+^) for a certain period of time. Protein from these four conditions is extracted and the expression of LC3-II is analyzed by Western blot, and normalized to reference proteins such as actin.

In the basal condition (EXP^−^), the amount of LC3-II in the absence of lysosomal inhibitors (BAF^−^) represents the APh pool in the steady state, analogous to taking a snapshot of the autophagic process (**Figure 2B**). The difference between the amount of LC3-II in cells incubated with and without lysosomal inhibitors (BAF^+^ – BAF^−^) in the basal condition represents the autophagosomes that have disappeared (i.e., the degradation phase), which is what is conventionally called autophagic flux. To calculate the autophagosomes that have formed, our model stems from the assumption that in the basal condition autophagy is at an equilibrium and therefore the size of the APh pool is constant (i.e., formation and degradation occur at the same speed):

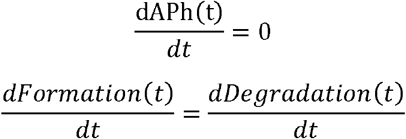

Thus, in the basal condition (EXP^−^) the autophagosomes that have formed are identical to the autophagosomes that have degraded, and thus are also represented by the amount of LC3-II with and without lysosomal inhibitors (BAF^+^ – BAF^−^).

In the experimental condition (EXP^+^), the amount of LC3-II in the absence of lysosomal inhibitors (BAF^−^ in EXP^+^) represents the size of the APh pool under the stimulus. Again, degradation can be calculated as the difference in LC3-II with and without lysosomal inhibitors [(BAF^+^ in EXP^+^) – (BAF^−^ in EXP^+^)]. Formation can be calculated as the difference between the amount of LC3-II in the presence of lysosomal inhibitors minus the size of the initial APh pool in steady-state conditions [(BAF^+^ in EXP^+^) - (BAF^−^ in EXP^−^)] (**Figure 2B, C**). This procedure allows us to calculate the formation and degradation of autophagosomes in control and experimental conditions (**Figure 2C, D**). To then compare whether the stimulus acts proportionally in both formation and degradation stages, we can calculate the ratio between experimental and basal conditions (EXP^+^/EXP^−^) for both formation and degradation, and the ratio between both as the net turnover ratio (**Figure 2C, E**). In basal conditions, the net turnover would be equal to one (red dotted line in **Figure 2E**).

### Dissecting out autophagosome formation and degradation

This model allows us to conceive all possible effects of the stimulus of interest on autophagy, as combinations of (increased/no change/decreased) autophagosome formation vs (increased/no change/decreased) autophagosome degradation (**Figure 3**). To better visualize the different possibilities (**Figure 3A**) we have numerically recreated seven possible scenarios (**Figure 3B-H**). A conventional stimulus of autophagy would be expected to increase the flux (i.e., degradation) and in addition our analysis would reveal that formation is similarly increased, leading to a net turnover ratio of one (**Figure 3B**). Therefore, under this stimulus autophagy increases but maintains an equilibrium: autophagosome formation and degradation increase proportionally, resulting in the maintenance of the net autophagic turnover but at a higher rate/velocity. This would be the expected case during treatment with the well-known autophagy activator rapamycin, an inhibitor of the mTORC1 complex (mechanistic target of rapamycin complex 1) (Civiletto et al., 2018). Similarly, the transcription factor-EB (TFEB) coordinately regulates the biogenesis of autophagosomes and lysosomes (Settembre et al., 2011) maintaining the equilibrium between formation and degradation (**Figure 4A**).

**Figure 3.**
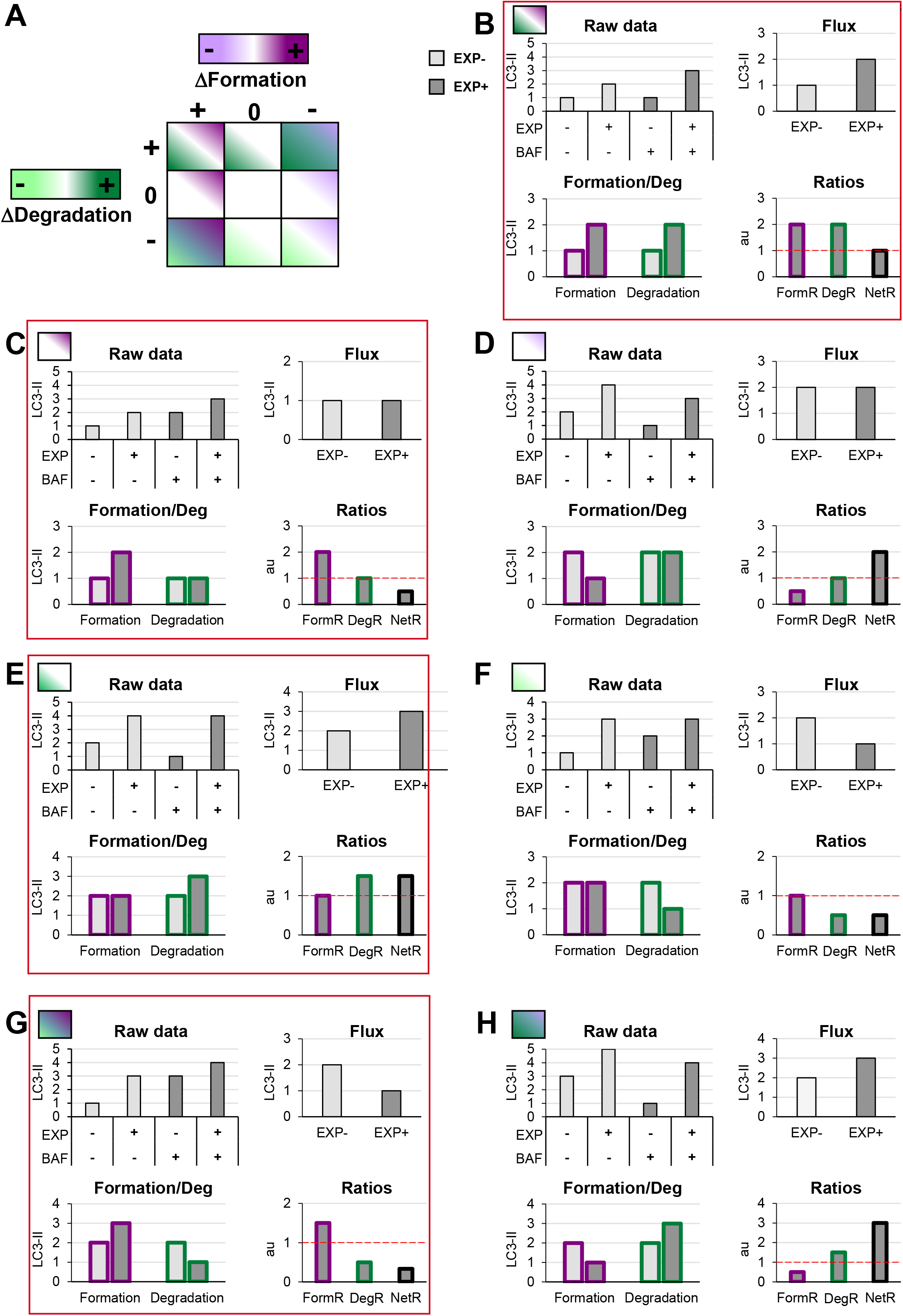
Combinatorial analysis of the effects of changes in autophagosome formation and degradation. **A**, Color-coded table representing changes in autophagosome formation (purple) and degradation (green). Dark tones represent increases, light tones decreases and white represents no changes. **B-H**, examples of analysis of autophagosome formation, degradation and net turnover from the table shown in A. The red dotted line represents the threshold of one to determine a significant change (over 1, basal conditions) in the formation, degradation and net turnover ratios. The panels underlined with red squares are explained in more detail with examples in **Figure 4**.

**Figure 4.**
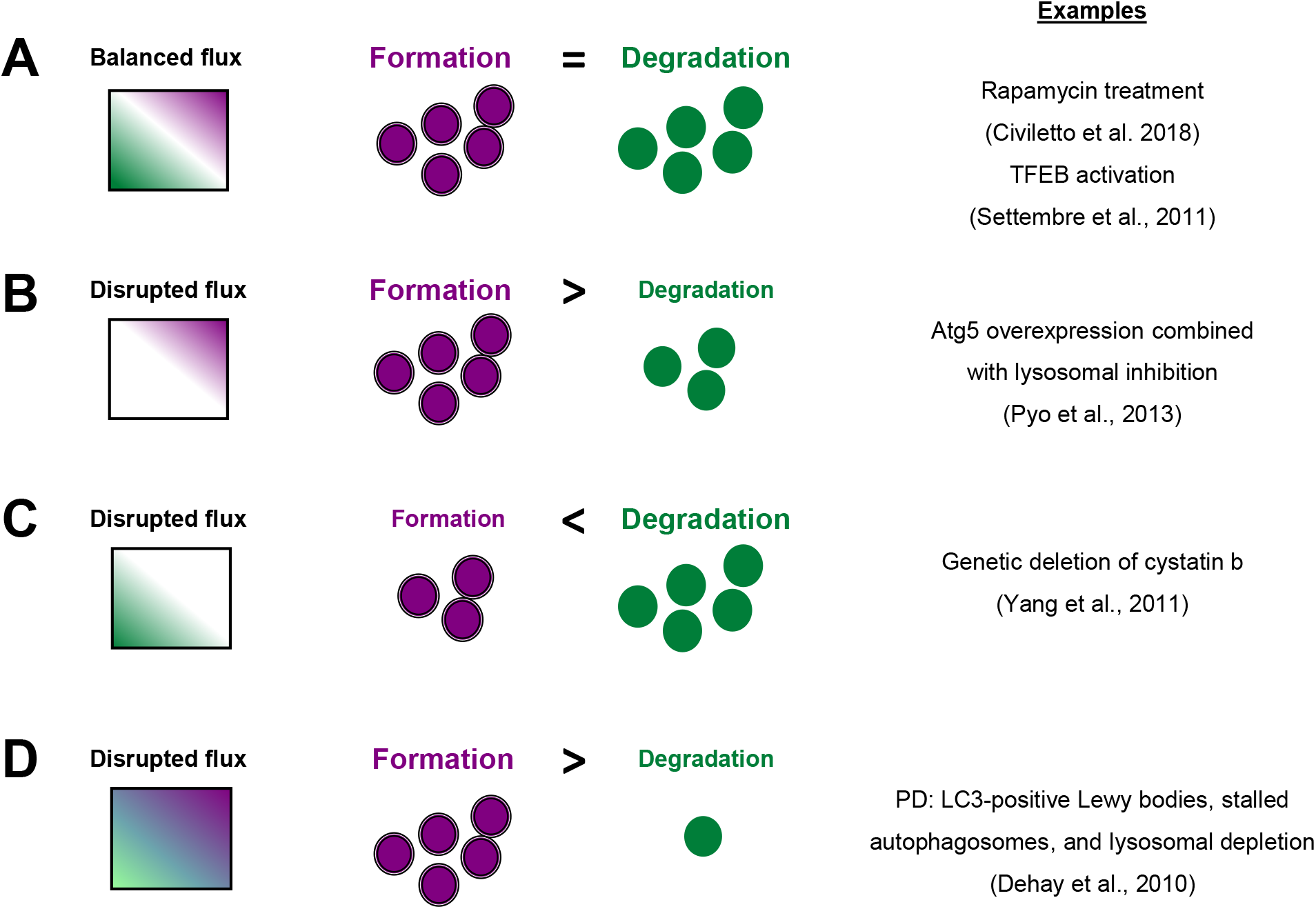
Experimental examples of variations in the formation and degradation of autophagosomes that lead to balanced or disrupted autophagy flux. **A**, After rapamycin treatment or TFEB translocation to the nucleus, autophagosome formation and degradation are proportionally increased, leading to a balanced up-regulation of autophagy flux. **B**, Atg5 overexpression elicits formation of autophagosomes. If lysosomes are inhibited under this scenario, autophagosomes are stalled and autophagy flux is disrupted. **C**, Genetic deletion of cystatin b, an enzymatic inhibitor of certain cathepsins, enhances cathepsin activity and lysosomal proteolysis, increasing degradation of autophagosomes without affecting formation, leading to disrupted autophagy flux. **D**, Parkinson’s disease (PD) dopaminergic neurons exhibit LC3-positive Lewy bodies, stalled autophagosomes and lysosomal depletion, which suggests dysfunctional flux due to increased formation and decreased degradation of autophagosomes.

In contrast, we can envision other conditions that differentially affect formation and degradation. For instance, some stimuli may selectively increase or decrease autophagosome formation without concomitantly affecting degradation (**Figure 3C, D**, respectively). As an example, overexpression of ATG proteins or accumulation of intracellular debris would lead to increased autophagosome formation. But if lysosomal efficiency (i.e., degradation) is not proportionally increased, autophagosomes will stall in the lysosomes without degrading the autophagic cargo, leading to a decreased net turnover ratio and increased autophagosome pool (**Figure 4B**). This effect has been for example observed in cells that overexpress Atg5 but whose lysosomal function is compromised (Pyo et al., 2013). In this case, calculation of the autophagy flux would not reveal any changes, concealing a potentially catastrophic situation for the cell that could not possibly be maintained over time.

It is also possible to imagine stimuli that selectively increase or decrease autophagosome degradation without affecting their formation (**Figure 3E, F**). For example, enhanced lysosomal biogenesis or lysosomal enzymes efficiency might lead to increased autophagosome degradation, resulting in an increased net turnover ratio and reduced autophagosome pool size (**Figure 4C**). This imbalance has been reported in mice genetically deficient for the cathepsin inhibitor cystatin B, which exhibit enhanced lysosomal proteolysis (Yang et al., 2011). Whereas in this case the calculation of the autophagy flux would suggest an enhanced autophagy, in fact cellular debris would not be removed any faster from the cytoplasm.

Finally, it is possible to encounter conditions in which formation and degradation are inversely regulated (**Figure 2G, H**). One possible case is a pathological scenario where dysfunctional organelles accumulate and the cell tries to enclose them in autophagosomes but lysosomal functionality is compromised, for instance because lysosomes are defective or engaged in other degradation pathways such as phagocytosis or endocytosis (**Figure 2G**). This effect could be observed in Parkinson’s disease (PD) dopaminergic neurons, which contain LC3-positive Lewy bodies, and have stalled autophagosomes, and lysosomal depletion (Dehay et al., 2010). This complex effect cannot be fully understood by simply analyzing the reduction in the autophagy flux but would be instead clearly described by our two-step model.

### Testing the model in vitro

We have directly validated our model with experimental data using two well-characterized autophagy modulators: the autophagy inducer rapamycin, which inhibits MTORC1 (Morel et al., 2017); and the autophagy inhibitor MRT68921, which blocks ULK1/2 (unc-51-like kinases 1/2) (Petherick et al., 2015; Morel et al., 2017). Both MTORC1 and ULK1/2 are early checkpoints of canonical autophagy: MTORC1 transduces signals from energy and damage sensors and is activated under stressful situations, releasing ULK1/2 (unc-51-like kinase 1/2) by phosphorylation to initiate the autophagy cascade (Plaza-Zabala et al., 2017; Mercer et al., 2018). As a cell model we used cultures of microglia (BV2 cell lines or primary cultures), the brain macrophages, and analyzed the amount of LC3-II by Western blot as a measurement of the size of the autophagosome pool.

In BV2 microglia rapamycin (6h, 100nM) showed the expected response and a tendency to increased LC3-II flux (**Figure 5A**). In addition, our model uncovered a parallel increase in formation and degradation of autophagosomes, resulting in a constant size of the APh pool and no changes in the net autophagosome turnover. Thus, rapamycin allowed the maintenance of the equilibrium between formation and degradation (**Figure 5B**), indicating a sustained autophagy that the cell can maintain over time.

**Figure 5.**
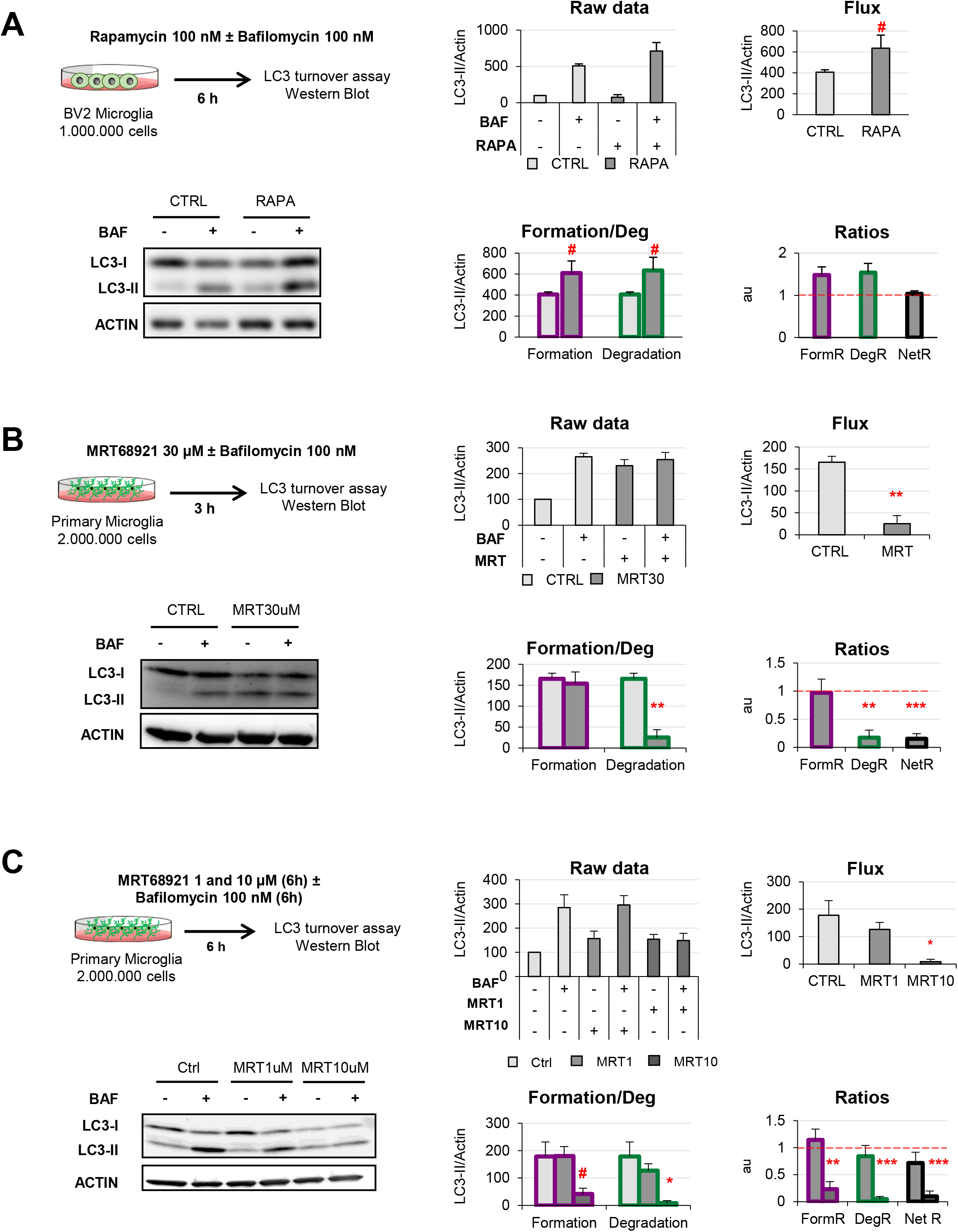
Validation of the two-step model with autophagy modulating compounds. **A**, Autophagy induction assessed after treatment with Rapamycin (100 nM, 6h) in the presence and absence of Bafilomycin (100 nM) in the BV2 microglia cell line. A representative blot, the raw data obtained, and the calculations of flux, autophagosome formation and degradation, and net turnover ratios are shown. **B, C**, Autophagy inhibition assessed after treatment with MRT68921 (30 μM, 3h in **B**; 1 and 10 μM, 6h in **C**) in the presence and absence of Bafilomycin (100 nM) in mouse primary microglia. A representative blot, the raw data obtained, and the calculations of flux, autophagosome formation and degradation, and net turnover ratios are shown. Data represent mean ± SEM of 3 independent experiments. # represents p<0.1, **\*** represents p<0.05 and ** represents p<0.01 by one tailed Student t-test (A, B), or Holm-Sidak after a significant effect of the treatment was found with 1-way ANOVA (C).

On the other hand, MRT68921 (3h, 30μM) resulted in the expected decrease in the LC3-II flux in primary microglia (**Figure 3B**). However, analysis with our model revealed that only degradation was reduced whereas autophagosome formation remained constant (**Figure 3B**). This data is in apparent contradiction with the described role of MRT in blocking the autophagy pre-initiation complex (Petherick et al., 2015; Morel et al., 2017). To address this discrepancy, we used a second paradigm of MRT68921 with a longer treatment and lower dosage (6h, 1-10μM; **Figure 5C**), and observed that the upstream effect of inhibition of autophagosome formation with MRT 10μM translated into a similar decrease in degradation (**Figure 5C**). Therefore, our model proves useful to discriminate the effect of experimental manipulations on the formation and/or degradation of autophagosomes.

### Future directions

Autophagy is a complex multi-step phenomenon and its assessment is a complicated task that requires using complimentary methods, as most current guidelines recommend (Mizushima et al., 2010; Klionsky et al., 2016). Visualization of double-membrane autophagosomes by transmission electron microscopy, live imaging of LC3 acidification using ratiometric analysis of fluorophores, or analysis of substrate degradation should corroborate the data obtained by analysis of LC3-II expression as a proxy for autophagosome formation and degradation. It is also important to note that autophagy is a time-dependent process and, as such, its dynamics should be assessed over time. In addition, LC3-II immunoblotting assays have several limitations, such as the reference protein used to normalize LC3-II values, the timing and concentration of the lysosomal inhibitor used, or the intrinsic nonlinear detection of proteins by enhanced chemoluminescence (ECL) (Rubinsztein et al., 2009). The most widely used method to assess autophagy, nonetheless, is the analysis of the LC3-II flux in the presence of lysosomal inhibitors. Nonetheless, the complexities associated to interpreting LC3-II flux have been thoroughly pointed out before, in the quest for an optimal “autophagomometer” (Rubinsztein et al., 2009). One of the key points is that autophagosomes formation and degradation are spatially and temporally dissociated and that therefore they need to be assessed independently.

To address this issue we here propose a simple conceptual frame to help interpreting LC3-II flux experiments. Our two-step model conceives the steady-state levels of LC3-II as an indirect measure of the pool of autophagosomes present when the snap-shot is taken. Assuming that in the basal condition the cells or tissue of interest are in some sort of equilibrium, the amount of autophagosomes formed and degraded should be roughly the same. Thus, the autophagosome pool can be treated as a black box to which the input (formation) and output (degradation) are identical, and can be estimated as the difference between LC3-II levels in the presence and absence of lysosomal inhibitors. In the experimental condition, degradation can be similarly calculated as the difference between LC3-II levels in the presence and absence of lysosomal inhibitors (i.e., the conventional LC3-II flux). In addition, we propose that the formation of autophagosomes in the experimental condition can be estimated by subtracting the steady-state autophagosome pool to the autophagosomes that have accumulated in the presence of lysosomal inhibitors. This model allows us to dissect out the effects of the experimental conditions to autophagosome formation and degradation. In addition, it also allows us to understand the net changes in the size of the autophagosomal pool that are the result of maintaining (or not) the net turnover ratio at equilibrium.

Nonetheless, our two-step model has several limitations that should be considered. The most important one is the assumption that autophagy (formation and degradation) are at equilibrium in the basal condition. This equilibrium implies coordinated control mechanisms that would be necessary to maintain autophagy in the long term term (Shen and Mizushima, 2014), but each cell type may have different regulation mechanisms under different metabolic constraints (Nwadike et al., 2018). Another important point is that autophagosome formation and degradation are not independent variables, as assumed in our model. For instance, it is obvious that if the lysosomal pool is not a limiting factor, the degradation will directly depend on the formation. In addition, feed-back mechanisms may link excessive lysosomal degradation with a subsequent reduction in autophagosome formation (Yu et al., 2010). In spite of these limitations, our model can provide a more expanded insight into the complexity of the autophagy process than simply analyzing the autophagic flux. In summary, we here show that using the LC3 turnover assay, our two-step model helps to systematically determine changes in autophagosome formation vs degradation, the net turnover and the size of the autophagosome pool to obtain a more comprehensive understanding of autophagy.

## METHODS

### Cell culture

The murine microglial BV2 cell line and primary microglia were used to test autophagy modulating compounds. BV2 microglia were grown and maintained in Dulbecco’s Modified Eagle Medium (DMEM) (Gibco) supplemented with Fetal Bovine Serum 10% (FBS, Gibco) and a mixture of antibiotics/antimycotic (1%) including, penicillin, streptomycin, and amphotericin (all from Gibco). For experiments, 1×10^6^ cells adhered to uncoated plastic plates were used. Primary microglia cultures were performed as previously described (Abiega et al., 2016; Beccari et al., 2018). Postnatal day 0-1 (P0-P1) fms-EGFP mice pup brains were extracted and the meninges were peeled off. The olfactory bulb and cerebellum were discarded and the rest of the brain was then mechanically homogenized by careful pipetting and enzymatically digested with papain (20U/ml, Sigma), and deoxyribonuclease (DNAse; 150U/μl, Invitrogen) for 15min at 37ºC. The resulting cell suspension was then filtered through a 40μm nylon cell strainer (Fisher) and transferred to a 50ml Falcon tube quenched by 5ml of 20% FBS (Gibco) in HBSS. Afterwards, the cell suspension was centrifuged at 200g for 5min, the pellet was resuspended in 1ml DMEM (Gibco) supplemented with 10% FBS and 1% Antibiotic/Antimycotic (Gibco), and seeded in T75 Poly L-Lysine-coated (15μl/ml, Sigma) culture flasks at a density of two brains per flask. Medium was changed the day after and then every 3–4 days, always enriched with granulocyte-macrophage colony stimulating factor (5ng/ml GM-CSF, Sigma). After confluence (at 37°C, 5% CO_2_ for approximately 14d), microglia cells were harvested by shaking at 100-150rpm, 37°C, 4h. Isolated cells were counted and plated at a density of 2×10^6^ cells/well on poly-l-lysine-coated plastic plates. BV2 and primary microglia were allowed to settle for at least 24h before experiments.

#### Drug treatments

BV2 microglia were treated with Rapamycin 100 nM (Fisher Scientific) for 6h in the presence and absence of Bafilomycin 100 nM (SelleckChem) for autophagy induction. Primary microglia were treated with the autophagy inhibitor MRT68921 1, 10 or 30 μM (Sigma) for 3 or 6h with or without Bafilomycin 100 nM (SelleckChem).

#### Protein extraction and Western Blot

Microglia were directly lysed in plastic plates with RIPA buffer containing protease inhibitor cocktail (100x) (ThermoFisher). The cell suspension was then sonicated for 5s and centrifuged (10,000g, 10min) to obtain solubilized protein in the supernatant. Sample protein content was quantified in triplicates by BCA (Bicinchoninic Acid) assay kit (ThermoFisher) at 590nm using a microplate reader (Synergy HT, BioTek). β-mercaptoethanol denatured proteins (15-20 ug) were loaded onto 14% Tris-glycine polyacrylamide gels (ThermoFisher) and run at 120V for 90min. Protein samples were then blotted to nitrocellulose membranes (0.45 μm pore size) (ThermoFisher) at 200 mA for 90min or using the Trans-Blot Turbo Mini Nitrocellulose Transfer Pack (Bio-Rad). Transfer efficiency was verified by Ponceau S (Sigma) staining. For immunoblotting, membranes were rinsed in Tris Buffered Saline containing 0.1% Tween 20 (Sigma) (TBS-T) and then blocked for 1h in TBS-T containing 5% powder milk. Membranes were afterwards incubated with rabbit primary antibody to LC3 (1:3000, NB100-2220, Novus Biologicals), and mouse primary antibody to β-actin (1:5000, Sigma), in TBS-T containing 4% Bovine Serum Albumin (BSA) overnight (4°C, shaker). Next day, membranes were rinsed and incubated with Horseradish Peroxidase (HRP) conjugated anti-rabbit (1:5000) and anti-mouse (1:5000) secondary antibodies (Cell Signaling) for the rapamycin blot or with the fluorescent StarBright Blue 700 anti-mouse (1:5000) and StarBright Blue 700 anti-rabbit (1:5000) secondary antibodies (Bio-Rad) for the MRT68921 blots in TBS-T containing 5% powder milk. After rinsing membranes, protein was visualized by enhanced chemiluminescence (ECL) using Supersignal West Femto Maximum Sensitivity Substrate (ThermoFisher) for the rapamycin blot or by immunofluorescence for the MRT68921 blots, in a ChemiDoc imaging system (BioRad). Band intensity was quantified using the Gel Analyzer method of Fiji software.

#### Statistics

Statistical analysis was performed with SigmaPlot. Normality and homoscedasticity were assessed prior to analysis. Raw LC3 data was initially analyzed by two-way ANOVA, but since an interaction between treatment (rapamycin, MRT68921) and bafilomycin was found, the global effect of the treatment was subsequently analyzed by one-way ANOVA. In addition, flux, and formation and degradation rates were analyzed by one-tail Student t-test (Figure 5A, B) or by one-way ANOVA followed by a Holm-Sidak posthoc test (Figure 5C). Formation and degradation rates were compared to one using a one-tail Student t-test. Data is shown as mean ± SEM. Only tests with p<0.05 are considered significantly different; tests with p<.1 are reported to have a tendency.

## Abbreviations

APh: Autophagosome
ATG: Autophagy-related protein
BAF: Bafilomycin A1
BCA: Bicinchoninic Acid
DegR: Degradation Ratio
DMEM: Dulbecco’s Modified Eagle Medium
ECL: Enhanced Chemoluminescence
EXP^−^: basal condition
EXP^+^: experimental condition
FBS: Fetal Bovine Serum
FormR: Formation Ratio
GM-CSF: Granulocyte-Macrophage Colony Stimulating Factor
LC3-II: microtubule-associated light chain protein 3 conjugated to phosphatidylethanolamine
mTORC1: Mechanistic Target of Rapamycin Complex 1
NetR: Net Ratio
PD: Parkinson’s disease
ss: steady-state
TBS-T: Tris Buffered Saline containing 0.1% Tween 20
TFEB: transcription factor-EB
ULK1/2: unc-51-like kinases 1/2

## ACKNOWLEDGEMENTS

This work was supported by grants from the Spanish Ministry of Science and Innovation (https://www.ciencia.gob.es/) with FEDER funds (RTI2018-099267-B-I00) and a Tatiana Foundation project (P-048-FTPGB 2018) to AS. VST holds a predoctoral fellowship from the Basque Government.

